# AspFlex: molecular tools to study gene expression and regulation in *Acinetobacter baumannii*

**DOI:** 10.1101/2023.03.17.533205

**Authors:** Merlin Brychcy, Alexis Kokodynski, Devin Lloyd, Veronica G. Godoy

## Abstract

*Acinetobacter baumannii* is a gram negative nosocomial opportunistic pathogen frequently found in hospital settings, causing high incidence of in-hospital infections. It belongs to the ESKAPE group of pathogens (the “A” stands for *A. baumannii*) that is known to easily develop antibiotic resistances. It is crucial to create a molecular toolkit to investigate its basic biology, such as gene regulation. Despite *A. baumannii* being a threat for almost two decades, an efficient and high-throughput plasmid system that can replicate in *A. baumannii* has not yet been developed. This study adapts an existing toolkit for *Escherichia coli* to meet *A. baumannii’s* unique requirements and expands it by constructing a mobile CRISPR interference (CRISPRi) system that can produce gene knockdowns in *A. baumannii*.

## Introduction

*Acinetobacter baumannii* is a worldwide nosocomial opportunistic pathogen with high morbidity and mortality^1^. *A. baumannii* infections have risen sharply during the last 20 years; the CDC classifies it as a serious threat level, accounting for 12,000 infections every year^2^*. A. baumannii’s* main strategies to be such a successful pathogen relies on acquisition of antibiotic resistances and ability to survive desiccation by existing in multicellular communities or biofilms^3^. Over half of the infections caused by *A. baumannii* are by multi drug resistant strains^4^.

Having a diverse array of molecular tools at our disposal is essential to understand the fundamental mechanisms governing basic molecular processes in *A. baumannii* to discover innovative strategies to eradicate it. Although fundamental tools and expression plasmids are already accessible^4,5^, such as plasmids for allelic exchange^6^ or transposon insertion^7^, they depend on conventional restriction enzyme cloning making them tedious, time consuming, inefficient, and the cloning of several inserts is cumbersome. More sophisticated genetic instruments, CRISPRi, are still in the early stages of development^5^. CRISPRi offers a valuable way to investigate essential genes in greater depth. This is especially significant considering the large number of genes of unknown function present in the *A. baumannii* genome. The main advantage of Golden Gate cloning is the use of type IIS restriction enzymes that cleave outside their recognition site. This enables the creation of homologous overhangs in a single or multiple fragments for directional cloning as well as combining digestion and ligation reactions in a single tube resulting in a significantly greater cloning efficiency^6,7^. This easy and efficient cloning system was a prime platform for the successful clone of gene editing fragments including TALEN^8^ and CRISPR/Cas9 sgRNAs^9,10^. Golden Gate cloning systems are well established for *E. coli*^11,12^, but they are currently unavailable for *A. baumannii*.

The aim of this study is to devise a Golden Gate based plasmid system for *A. baumannii*, which will facilitate the simple and extremely efficient insertion of recombinant fragments of interest into plasmids. We report the creation of a plasmid kit (AddgeneIDs: 1000000217) with the possibility to express up to 20 transcriptional units at once by adjusting the already established EcoFlex^11^ system to *A. baumannii’s* needs. We demonstrate here its use by creating an *A. baumannii* mobilizable CRISPRi gene system.

## Results

### Construction of an AspFlex cloning system for *A. baumannii*

The EcoFlex^11^ system is organized in a standard modular cloning (MoClo) fashion based on multiple levels of organization. Level 0 plasmids contain elements such as promoters, terminators, or open-reading frames. These can be used to build a transcriptional unit (TU). Level 1 plasmids allow for the expression of a single TU by combining level 0 elements. Subsequently, level 2 plasmids allow the expression of multiple TUs by combination multiple level 1 plasmids and so on.

To adapt EcoFlex^11^ to *A. baumannii* a specific set of plasmid replication elements is needed. We introduced the replication elements from pWH1266 a plasmid from *A. calcoaceticus^1^* known to replicate in *A. baumannii*^14^, but also other *Acinetobacter spp*^15^. The replication elements were amplified using primers ori_ab_psti_f and ori_ab_psti_r (Supplemental table 1) and introduced into the level 1-3 plasmids using PstI-HF (New England Biolabs) followed by ligation with T4 DNA ligase (New England Biolabs). Since *A. baumannii* is resistant to Chloramphenicol^16–18^, the level 2 plasmid *cat* gene encoding a chloramphenicol acetyltransferase resistance cassette was digested with EcoRV-HF (New England Biolabs), and a blunt ended fragment containing the *kanR* gene fragment with its own promoter was inserted resulting in interruption of *cat*.

The result of these manipulations is a set of level 1 (pMB1-A through pMB1-E; AddgeneID: 190114-190119), level 2 (pMB2a through pMB2-D; AddgeneID: 190120-190125) and level 3 (pMB3-A and pMB3-B; AddgeneID: 190126 and 120127) plasmids. To test whether plasmids would stably replicate and be maintained in *A. baumannii*, we used a pMB1-A derivate, a level 1 plasmid strongly expressing *egfp*^19^ and introduced it by transformation into *A. baumannii*. We expected that *A. baumannii* cells in which the plasmid was successfully replicated would be both resistant to the plasmid marker and brightly green fluorescent for multiple generations. This is indeed what we found (see Supplemental Figure 1).

### AspFlex and CRISPRi

To create the mobilizable CRISPRi gene system we first produced the dCas9 plasmid. To do so elements such as an ATc inducible promoter (pTet) with the pET-RBS ribosomal binding site (AddgeneID: 72981) to tightly regulate the expression of the catalytically deficient Cas9 gene from pdCas9-bacteria (AddgeneID: 44249)^20^, and the Bba_B0012 terminator (AddgeneID: 190129) were moved onto pMB1-A. The catalytically deficient Cas9 gene^20^, *dCas9*, was amplified using primers pdCas9_new_r and pdCas9_new_f (Supplemental table 3). The resulting fragment was cloned into a level 0 plasmid. However, to move this fragment to the pMB1-A plasmid on AspFlex, a BsmBI site, which is a type II restriction enzyme required for cloning, needed to be removed from the dCas9 fragment. To achieve this, site-directed mutagenesis was carried out with the primers pBP_cas9_mut_r and pBP_cas9_mut_f (Supplemental table 3).

A second plasmid carrying the guide RNAs (sgRNAs) also needed to be implemented. For easy selection and Golden Gate compatible cloning of sgRNAs, we designed a custom fragment (Figure 1). Here, a *lacZ* alpha fragment for blue-white screening is flanked on one side by the strong promoter J23119^5^ and on the other by sequences encoding the tracrRNA (Figure 1 (A)). A simple reaction with two complimentary oligonucleotides including the Bpi-HF (NEB) sequence plus two distinct overhangs were used to generate directional cloning of a functional sgRNA(Fig. 1A). Thus, the AspFlex plasmid, pBP_sgRNA, (AddgeneID: 190128) allows easy cloning of sgRNAs (Figure 1 (B)). Screening of white positive colonies on X-Gal plates is carried out by PCR with flanking primers. We determined cloning efficiency by counting white colonies on X-Gal that contained the expected construct and found this is excellent; 100% of white colonies analyzed were correct (see Supplemental Figure 3).

**Figure 1:**
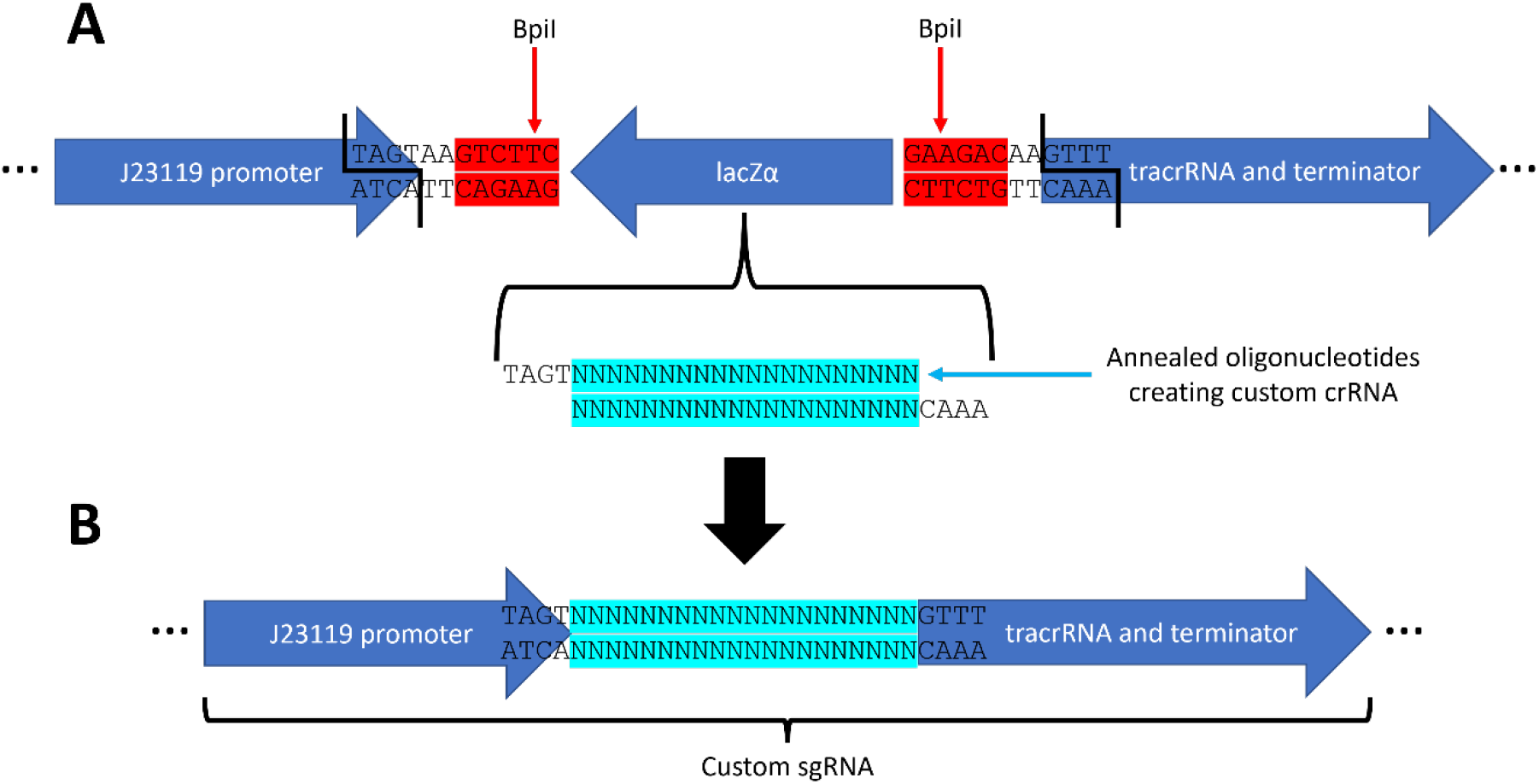
Schematic of level 0 fragment included for cloning into the pBP_sgRNA plasmid. (A) The pBP_sgRNA plasmid contains the promoter, tracrRNA and terminator elements needed for the expression of a sgRNA. Additionally, it contains a *lacZ* alpha fragment flanked by BpiI, a type II restriction enzyme, recognition sites. Because BPiI digests away from its recognition site, it was possible to create custom overhangs within the promoter/tracrRNA fragments to create a directional and seamless expression unit. Shown underneath the diagram is a representation of a custom CRISPR RNA (crRNA) that would be created by annealing two complimentary oligonucleotides. (B) Depiction of the resulting guide RNA (sgRNA) using BpiI mediated cloning in which the *lacZ* gene is replaced with the custom crRNA generating a ready to use sgRNA expression unit which will be present in white colonies on X-Gal plates.

To test the CRISPRi system in *A. baumannii*, two level 2 proof-of-principle plasmids were constructed. One of these plasmids, pMB04, contains an ATc inducible *dCas9* gene, a strongly expressed *egfp* and *mcherry* and sgRNAs targeting the fluorescence protein genes. The other, pMB05, contains the same fluorescence genes and a no-targeting nonsense sgRNAs (see supplemental information for sequence of both plasmids). We tested fluorescence intensity of *A. baumannii* ATCC17978 strains with either plasmid with different concentrations of ATc and without ATc (Fig. 2). We found that in ATc treated cells with pMB04, the plasmid with the fluorescent proteins targeting sgRNAs, the fluorescence signal is significantly repressed, but not with pMB05 carrying the nonsense non-targeting sgRNA (see Figure 2 A-D). We found that increasing concentrations of ATc intensified this effect. The data indicate that the CRISPRi platform is effective in *A. baumannii*. Furthermore, the platform also allows *dCas9* expression with different promoters, constitutive or inducible (for example pBAD, included in the set; AddgeneID: 190132).

**Figure 2:**
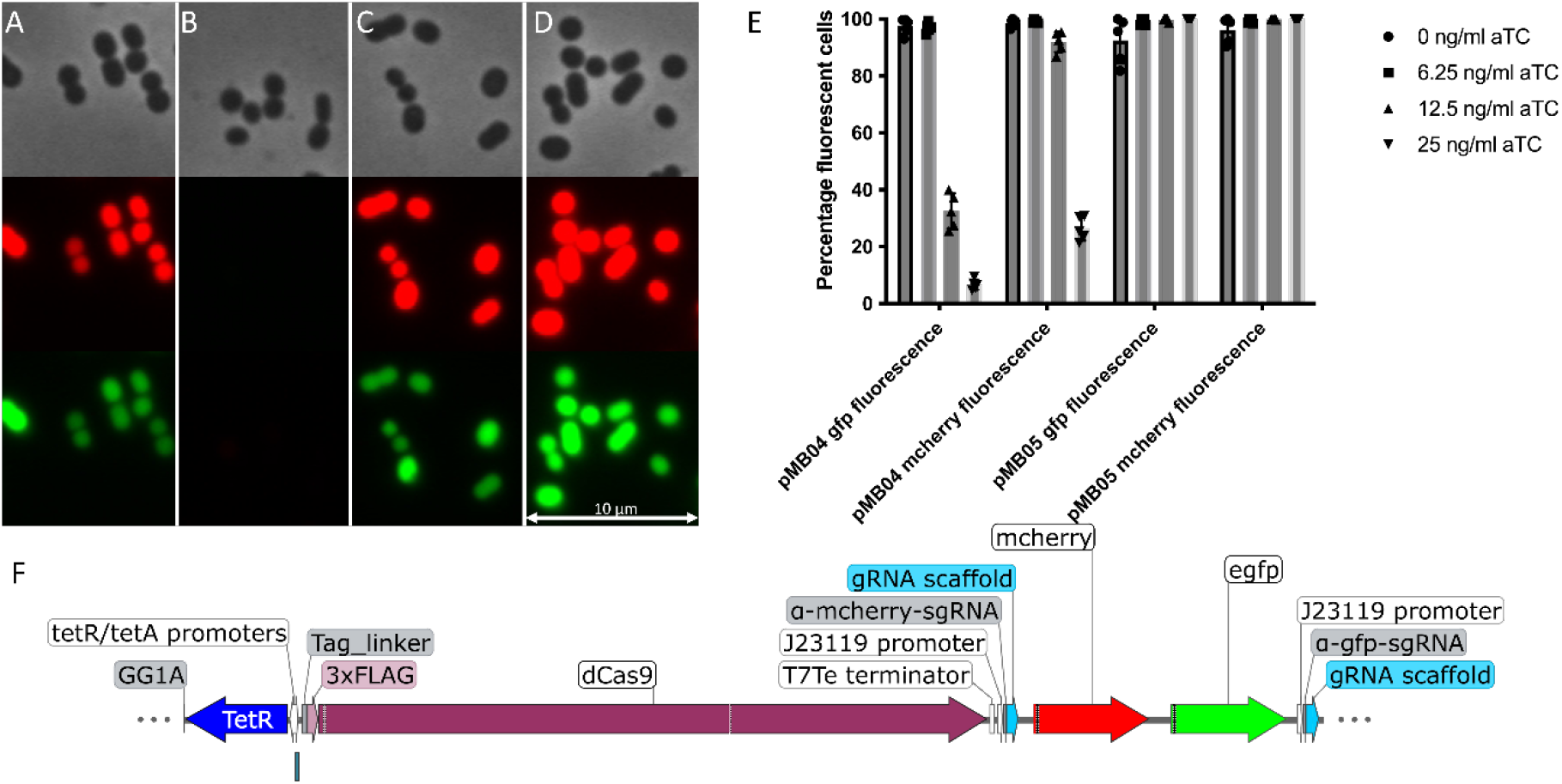
Fluorescence is suppressed by CRISPRi. (A)-(D) show phase contrast microscope images in the top, red fluorescence channel microscope image in the middle and green fluorescence channel in the bottom. Data was collected by subculturing saturated cells with ATc for 3h, with subsequent microscopy analysis and quantification of fluorescence with ImageJ^21^/Fiji^22^. (A) *A. baumannii* containing pMB04 construct (sgRNAs targeting fluorescence genes) with no ATc, the *dCas9* gene inducing agent. (B) *A. baumannii* containing pMB04 construct with 25 ng/mL of ATc. (C) *A. baumannii* with pMB05 construct (nonsense sgRNAs) with no ATc. (D) *A. baumannii* cells with pMB05 with 25 ng/mL ATc. (E) Quantification of percentage of fluorescent cells at different concentrations of inducing agent ATc. The percentage of fluorescent cells is significantly reduced only when sgRNAs targeting the fluorescence genes are present and not observed when nonsense sgRNAs are used. (F) Insert of plasmid pMB04. The insert contains an ATc inducible dCas9 (blue and plum), two fluorescence genes (egfp and Dsred; green and red) as well as sgRNAs targeting the two (grey and teal) genes. The binding site of the sgRNAs on the appropriate fluorescence gene is marked with a black line within the fluorescence gene. Image was generated with SnapGene (http://www.snapgene.com).

The obtained results from these experiments demonstrate the effective use of a Mo-Clo-based CRISPRi plasmid system in *A. baumannii*, providing opportunities for further research on essential genes and gene regulation. By using this system, it is possible to precisely control the expression of a regulator and evaluate its impact. This approach presents a wide range of possibilities for studying *A. baumannii*.

In summary, AspFlex shows to be an ideal, efficient, fast, and easy method to construct a wide range of genetic tools such as transcriptional reporters, or protein fusions. We demonstrate here its use by setting up a mobilizable CRISPRi system that allows the regulation of multiple transcripts at once, providing a wide range of possibilities to study the functionality of gene pathways in *A. baumannii*.

## Experimental procedures

Detailed protocols for all methods used in this report are in the supplemental information.

### Design of sgRNA oligos and cloning in level 0 plasmids

To create a CRISPRi sgRNA, appropriate sites in the promoter region/early open reading frame of the fluorescent genes containing an NGG PAM site were identified. Two complementary oligonucleotides of 20-24 bp length were designed with the one on the same strand as the NGG having a TAGT 5’ and the complementary oligonucleotide having a AAAC 5’ overhang (see Supplemental Table 3). The two oligonucleotides were annealed in ligase buffer by heating to 95 °C for 5 minutes and subsequent incubation at 22 °C for 20 minutes. A Golden-Gate reaction using BpiI, T4-DNA ligase, pBP_sgRNA (AddgeneID: 190128) and the annealed oligos was then used to clone the custom sgRNA. Positive clones were screened using Blue-White color formation in plates with X-Gal. The recombinant plasmid can be used in subsequent Golden Gate reactions to construct a level 1 MoClo plasmid, with a level 1 target plasmid (AddgeneID: 190114-190119), BsaI, and T4-DNA Ligase (both NEB). In level 1 and 2 plasmids the cloning site for the ORF disrupts the mCherry gene, interfering with its expression such that the successful integration of an ORF would result in the disappearance of *mCherry* and a color shift from red to white.

### Cloning of CRISPRi expression units

To create a full CRISPRi expression unit, the level 1 plasmids containing *dCas9* and the sgRNAs need to be combined in a level 2 plasmid. For this, a level 2 Golden Gate reaction using the appropriate plasmids (AddgeneIDs: 190120-190125), BsmBI and T4-DNA ligase and the previously generated level 1 plasmids were used. Positive clones were screened via Red-White colony colors as indicated above. The system allows the expression of up to 20 sgRNAs at the same time.

## Abbreviations

ESKAPE: *<u>E</u>nterococcus faecium; <u>S</u>taphylococcus aureus; <u>K</u>lebsiella pneumoniae; <u>A</u>cinetobacter baumannii; <u>P</u>seudomonas aeruginosa; <u>E</u>nterobacter spp* 6 organisms of critical importance in inhospital infections because of their abilities to gain antibiotic resistances.
MoClo: <u>M</u>odular <u>Clo</u>ning. Plasmids sets specifically created for Golden Gate cloning.
CRISPRi: <u>C</u>lustered <u>r</u>egularly <u>i</u>nterspaced <u>s</u>hort <u>p</u>alindromic <u>r</u>epeats <u>i</u>nhibition. An expression inhibition system based on a deactivated Cas9 protein physically blocking transcription of a genomic region through an sgRNA binding in that region.
egfp: <u>e</u>nhanced green <u>f</u>luorescent <u>p</u>rotein.
ATc: <u>A</u>nhydrous <u>T</u>etracycline
X-Ga1: 5-bromo-4-chloro-3-indolyl-β-D-galactopyranoside. An organic compound used for blue-white screening.

## Acknowledgements

V.G-C. is funded by a stipend from Northeastern University Skills for Capacity and Inclusion, an Inclusive Excellence award from HHMI.

The EcoFlex kit was a gift from Paul Freemont (Addgene kit #1000000080) and pdCas9-bacteria was a gift from Stanley Qi (Addgene plasmid # 44249; http://n2t.net/addgene:44249; RRID:Addgene_44249).

We would like to thank the Geisinger Lab at Northeastern University for their help in developing a CRISPRi platform for *A. baumannii* and building the framework for it.

We would also like to thank the Chai Lab at Northeastern University for their support, feedback, and equipment.

