## Supplemental Information for "AspFlex: molecular tools to study gene expression and regulation in *Acinetobacter baumannii*"

**Bacterial strains and growth conditions**

*Escherichia coli* DH5α was used for all initial molecular cloning. *A. baumannii* ATCC 17978 was prepared for electroporation to introduce the chosen plasmids[1]. Both strains were grown in LB Miller (10 g/L NaCl; 10 g/L Tryptone; 5 g/L yeast extract; 20 g/L Agar (for plates)). Selection was performed through addition of necessary antibiotics; Carbenicillin 100 µg/mL and Kanamycin 35 µg/mL in *A. baumannii* and Chloramphenicol 20 µg/mL for *E. coli*. Strains were grown at 37°C at 225 rpm for 16 h.

**Microscope imaging and fluorescence quantification**

Single cell fluorescence microscopy was carried out to quantify fluorescence in strains with the CRISPRi test plasmids pMB04 (targeting sgRNA) and pMB05 (non-targeting sgRNA). Images were quantified with ImageJ/Fiji[2]. The fluorescence intensity was measured in 10 sets of microscope images with at least 500 cells per set. The number of fluorescing cells was divided by the total number of cells to obtain the percentage of fluorescent cells[2].

To prepare the cells for imaging a saturated culture of the appropriate *A. baumannii* strain containing the plasmid to be tested was grown for at least 16h at 37°C in LB medium with antibiotic (in this case Kanamycin). The saturated culture was diluted 1:1000 into fresh LB medium (no antibiotic added) with the appropriate concentration of ATc, the inducing agent, until reaching exponential phase (around 3 h). 2 μL of bacterial cells were then deposited on 1% agarose pads on microscope slides[3]. Cells were imaged in a Leica DM5000 DFC3000 G microscope with a 100x magnification. Images were taken in phase contrast, with a Texas red filter (Excitation: 540 nm – 580 nm; Dichromatic mirror: 595 nm; Emission: 607 nm – 683 nm) or a gfp filter (Excitation: 450 nm – 490 nm; Dichromatic mirror: 495 nm; Emission: 500 nm – 550 nm) (See Figure 2).

**Design and cloning of sgRNAs for CRISPRi**

The process of generating a complete CRISPRi construct involves multiple steps using AspFlex. The initial stage requires identifying a potential sgRNA site, which is ideally located within the promoter region or close to the 5’ end of the open reading frame of the protein target intended for CRISPRi-knockdown. While sgRNAs that are complementary to the non-template strand are preferred, we have also observed effective outcomes with sgRNAs targeting the template strand. Once the appropriate NGG PAM site is identified in the regions mentioned above, the next step involves designing two complementary oligonucleotides specific to that sgRNA. The oligonucleotide situated on the same strand as the NGG site should possess a TAGT 5’ overhang, while the complementary oligonucleotide needs an AAAC 5’ overhang (Supplemental Figure 3).

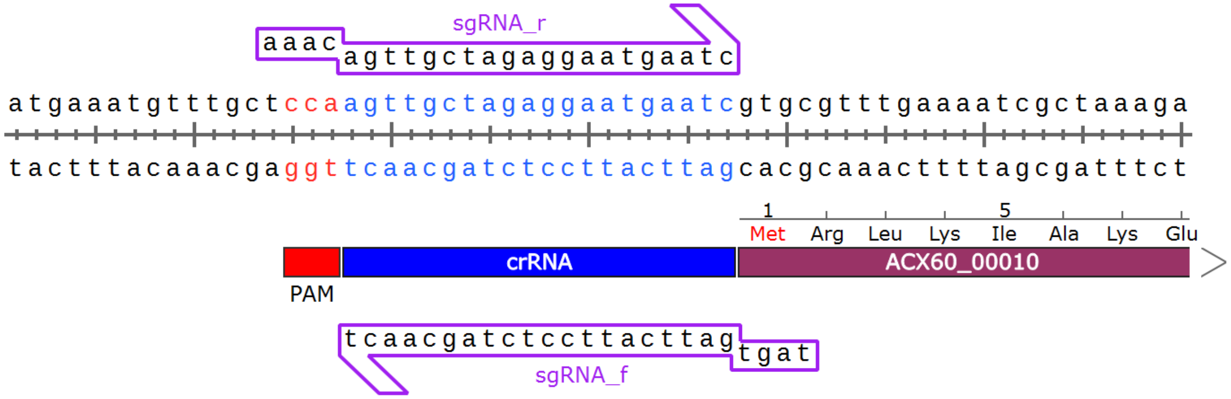


Supplemental Figure 3: Example of design of an sgRNA for CRISPRi. The crRNA is shown in blue, PAM in red. The gene suppressed by CRISPRi is shown in plum and the two complimentary oligonucleotides are shown in purple. Figure generated with Snapgene (http://snapgene.com).

The two oligonucleotides were annealed to create double stranded DNA with the necessary overhangs for the Golden Gate reaction. For this, we combined 5 µL of 10x T4 DNA ligase buffer (NEB) with 5 µL of each of the oligonucleotides at 100 µM in a PCR reaction tube. The PCR reaction tube was incubated in a Thermocycler at 95 °C for 5 minutes, followed by 22 °C for 20 minutes.

A Golden Gate reaction with the annealed oligonucleotides was then set up. The reaction had a final volume of 10 µL and contained 1 µL of T4 DNA ligase buffer (NEB), 1 µL of the annealed oligonucleotides, 0.03 pmol (~ 50 ng) of the pBP_sgRNA plasmid, 0.25 µL of BpiI-HF (NEB) and 0.5 µL of T4 DNA ligase (NEB). The reaction mix was incubated in the thermocycler with the following settings: Cycling 50 times with a first step at 37 °C (digestion step) for 2 minutes, followed by 16 °C for 5 minutes (ligation step). Afterwards, the sample was incubated for 1 hour at 37 °C to digest left-over backbone plasmid, followed by an 80 °C step for 20 minutes to heat-inactivate the enzymes.

The reaction mixture is then introduced by transformation into *E. coli* competent cells (for example DH5α) and deposited on LB medium plates containing Chloramphenicol, as well as IPTG and X-Gal for blue-white screening. White colonies are confirmed via colony PCR the next day.

While the reactions are generally highly efficient and correct, we still recommend sequencing all plasmids to avoid SNPs. Positive clones yield a successful level 0 sgRNA construct.

Afterwards, a level 1 plasmid was created with the successfully constructed level 0 sgRNA plasmid. The backbone selected in this case depends on the amount of sgRNAs expressed (see supplemental table 2). Considering that the *dCas9* constructs are present in pMB1-A plasmids, sgRNAs should never be cloned into pMB1-A, because multiple pMB1-A constructs would lead to incompatible overhangs in level 2 plasmids. For a single sgRNA construct, the guide RNA needs to be cloned into pMB1-B. For this in a 10 µL reaction, 30 pmols (~150 ng) of pMB1-B (or accordingly to other level 1 plasmid) with 0.03 pmol (50 ng) of the level 0 construct, 1 µL of T4 DNA ligase buffer, 0.25 µL of BsaI-HFv2 (NEB) and 0.5 µL of T4-DNA ligase (NEB) were combined. Mixes were incubated with the same settings as the level 0 reaction, introduce by transformation into standard *E. coli* competent cells (for example DH5α) and deposited on LB medium plates containing Ampicillin/Carbenicillin as a selection agent. The successful clones are white and were confirmed with Colony-PCR and sequencing.

To finish the cloning process, a level 2 plasmid containing both the sgRNAs and the *dCas9* needed to be created. In the case of a single guide RNA, in a 10 µL reaction, 1 µL of T4-DNA ligase buffer needs to be combined with 0.015 pmol (75 ng) of both of the level 1 constructs (in this case the sgRNA in pMB1-B and the *dCas9* in pMB1-A), 0.015 pmol (75 ng) of the level 2 backbone (in this case pMB2a; see supplemental table 2) and 1 µL of NEBridge Golden Gate assembly BsmBI-HFv2 master mix. Mixtures were incubated in the thermocycler with the following settings: 50 cycles at 42 °C for 5 minutes first, then at 16 °C for 5 minutes. Enzymes were heat -inactivated with a cycle at 60 °C for 5 minutes.

The reaction mix was then introduced by transformation into competent *E. coli* cells (we recommend highly competent commercial cells, for example NEB® 10-beta) selecting on LB medium with Kanamycin and screening for white colonies. Colony PCR and sequencing are recommended. A control digest, considering the size of the plasmid, could also be performed. The constructs can be introduced by transformation into *A. baumannii* once the sequence is confirmed.

If a level 3 plasmid needs to be created, follow the instructions for a level 1 plasmid.

To induce CRISPRi the optimal concentration of ATc required depends on the level of suppression needed of the protein targeted for knock-down. Generally, a full CRISPRi knockdown can be observed by adding 200 ng/mL of ATc on plates. However, in liquid culture, lower concentrations of ATc (for example 50 ng/mL) can be used. Our recommendation is to first grow a saturated overnight culture with the selective agent (Kanamycin) and then outgrow the cells with and without ATc until they reach exponential phase. We do not advise to use the selective agent (Kanamycin) when activating expression of the *dCas9* gene. Plasmid stability is insured via a toxin-antitoxin system on the plasmid guaranteeing replication of the plasmid even without a selecting agent. The successful knock-down can be detected either via qPCR or through phenotypical changes associated with the knock-down.

**Example: Cloning of a transcriptional reporter with AspFlex and EcoFlex**

To yield the full power of AspFlex, we heavily recommend acquiring the EcoFlex kit by the Freemont lab (Addgene kit #1000000080). This kit will provide additional plasmids expanding the possibilities the AspFlex kit offers, especially regarding expression or reporter plasmids. To create a transcriptional reporter with AspFlex, multiple level 0 elements from EcoFlex can be used. For example, the pET-RBS (Addgene Nr. 72981), the *egfp* gene (Addgene Nr. 72960) and the Bba_B0012 terminator (Addgene Nr. 72997). The promoter region is amplified with primers containing

BsaI sites and the according overhangs (CTAT and GTAC). A Golden Gate level 1 reaction like the one described above (use BsaI as restriction and combine all the elements mentioned above with a level 1 plasmid of choice) can be set up and introduced by transformation into *E. coli.* The plasmids can be introduced by transformation into *A. baumannii* to serve as transcriptional reporters after confirmation via colony PCR and sequencing.

**Oligonucleotides**

Supplemental Table 1: Oligonucleotides used in this work.

| Oligonucleotide name | Sequence (5' - 3') |
|---|---|
| ori_ab_pstI_r | AAATTCTGCAGTTAACAAGTTGCCTGACGCC |
| ori_ab_pstI_f | ATATTCTGCAGTTGCAAGACAATATCGACCG |
| pCas9_new_r | attggtctcttcgagttagtcacctcctagctgac |
| pCas9_new_f | attGGTCTCacataatggataagaaatactcaataggcttag |
| pBP_cas9_mut_r | ttccgctgtctctccactgt |
| pBP_cas9_mut_f | gcgactcgtcttaaacggac |
| mCherry_sgRNA_f | tagtcaagggcgaggaggataaca |
| mCherry_sgRNA_r | aaactgttatcctcctcgcccttg |
| GFP_sgRNA_r | aaacgtgaaaagttcttctccttt |
| GFP_sgRNA_f | tagtaaaggagaagaacttttcac |
| nonsense_sgRNA_r | aaacCTGCCATACCAGGCGCGTAC |
| nonsense_sgRNA_f | tagtGTACGCGCCTGGTATGGCAG |
| seq_primer | atttcagataaaaaaaatccttagctttcg |

#### Plasmids

Supplemental Table 2: Plasmids used in this work.

| Plasmid name | Level | Purpose | Antibiotic marker | Selection marker | Restriction enzyme | Source |
|---|---|---|---|---|---|---|
| pMB1-A | 1 | Cloning of single TU | Carbenicllin | *rfp* | BsaI | This study |
| pMB1-B | 1 | Cloning of single TU | Carbenicllin | *rfp* | BsaI | This study |
| pMB1-C | 1 | Cloning of single TU | Carbenicllin | *rfp* | BsaI | This study |
| pMB1-D | 1 | Cloning of single TU | Carbenicllin | *rfp* | BsaI | This study |
| pMB1-D1 | 1 | Cloning of single TU | Carbenicllin | *rfp* | BsaI | This study |
| pMB1-E | 1 | Cloning of single TU | Carbenicllin | *rfp* | BsaI | This study |
| pMB2-a | 2 | Cloning of 2 Tus | Kanamycin | *rfp* | BsmBI | This study |
| pMB2-b | 2 | Cloning of 3 Tus | Kanamycin | *rfp* | BsmBI | This study |
| pMB2-A | 2 | Cloning of 4 or 5 TUs | Kanamycin | *rfp* | BsmBI | This study |
| pMB2-B | 2 | Cloning of 4 or 5 TUs | Kanamycin | *rfp* | BsmBI | This study |
| pMB2-C | 2 | Cloning of 4 or 5 TUs | Kanamycin | *rfp* | BsmBI | This study |
| pMB2-D | 2 | Cloning of 4 or 5 TUs | Kanamycin | *rfp* | BsmBI | This study |
| pMB3-A | 3 | Cloning of 2 fragments from level 2 | Carbenicllin | *rfp* | BsaI | This study |
| pMB3-B | 3 | Cloning of 4 fragments from level 2 | Carbenicllin | *rfp* | BsaI | This study |
| pBP_sgRNA | 0 | Cloning of level 0 sgRNA | Chloramphenicol | *lacZ* | BpiI | This study |
| pMB1-A-ptet-dCas9 | 1 | Level 1 plasmid expressing ATc inducible dCas9 | Carbenicllin | | BsmBI | This study |
| pMB1-A-ptet-FLAG-dCas9 | 1 | Level 1 plasmid expressing ATc inducible FLAG tagged dCas9 | Carbenicllin | | BsmBI | This study |
| pBP_dCas9 | 0 | Level 0 plasmid containing point mutated dCas9 | Chloramphenicol | | BsaI | This study |
| pMB04 | 2 | Proof of principle sgRNAs targeting the fluorescence proteins | Kanamycin | | | This study |

| | | | | | | |
|---|---|---|---|---|---|---|
| pMB05 | 2 | Proof of priniciple, nonsense sgRNAs | Kanamycin | | | This study |

Supplemental Table 3: Table showing which plasmids are needed to create plasmids with certain amounts of transcriptional units.

| Amount of TUs | Plasmids needed | Alternative plasmids (same level) |
|---|---|---|
| 1 | pMB1-A | pMB1-B; pMB1-C; pMB1-D; pMB1-D1; pMB1-E |
| 2 | pMB1-A, pMB1-B, pMB2-a | |
| 3 | pMB1-A, pMB1-B, pMB1-C, pMB2-b | |
| 4 | pMB1-A, pMB1-B, pMB1-C, pMB1-D, pMB2-A | pMB2-B; pMB2-C; pMB2-D |
| 5 | pMB1-A, pMB1-B, pMB1-C, pMB1-D1, pMB1-E, pMB2-A | pMB2-B; pMB2-C; pMB2-D |
| 6 | pMB1-A, pMB1-B, pMB2-a, pMB1-A, pMB1-B, pMB1-C, pMB1-D, pMB-2B, pMB3-A | |
| 7 | pMB1-A, pMB1-B, pMB2-a, pMB1-A, pMB1-B, pMB1-C, pMB1-D1, pMB1-E, pMB2-B, pMB3-A | |
| 8 | pMB1-A, pMB1-B, pMB1-C, pMB1-D, pMB2-A, pMB1-A, pMB1-B, pMB1-C, pMB1-D, pMB2-B, pMB3-A | |
| 9 | pMB1-A, pMB1-B, pMB1-C, pMB1-D1, pMB1-E, pMB2-A, pMB1-A, pMB1-B, pMB1-C, pMB1-D, pMB2-B, pMB3-A | |
| 10 | pMB1-A, pMB1-B, pMB1-C, pMB1-D1, pMB1-E, pMB2-A, pMB1-A, pMB1-B, pMB1-C, pMB1-D1, pMB1-E, pMB2-B, pMB3-A | |
| | See EcoFlex Kit manual for further guidance with more fragments. | |

### Supplemental Figures

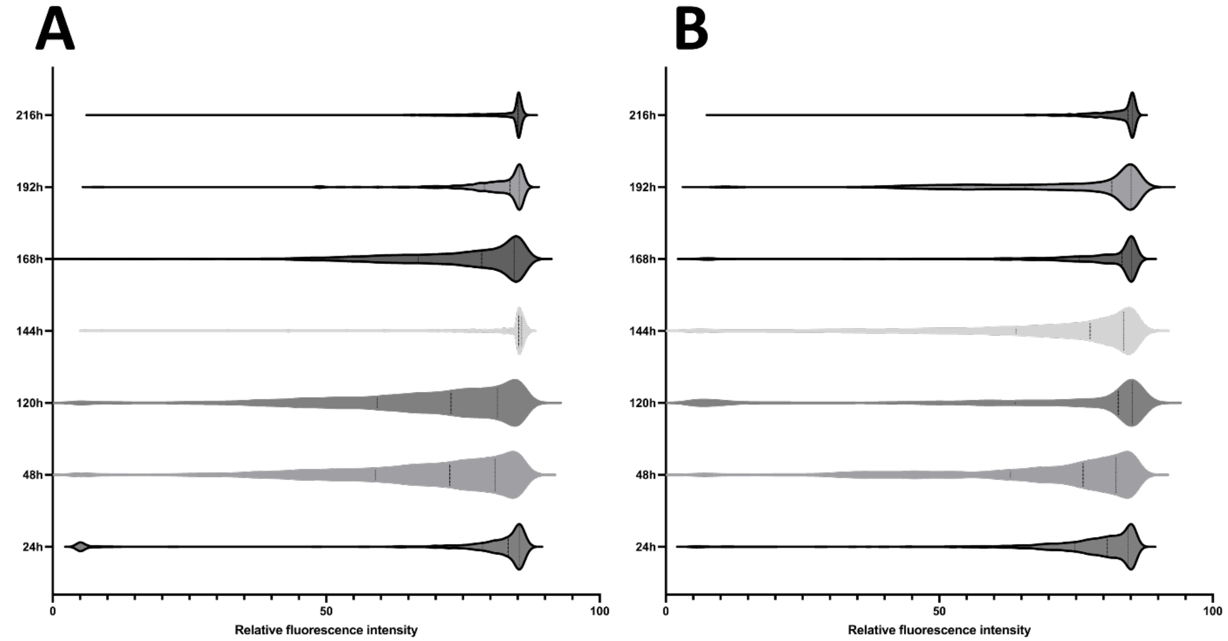


Supplemental Figure 1: AspFlex plasmids show stable replication even after multiple generations. The plasmid pMB1-A (AddgeneID: 190114) containing a strongly expressed *egfp* was introduced by transformation into *A. baumannii* ATCC17978. Plasmid stability was determined by diluting the culture 1:1000 dilution every 24h for 216h in total and measuring the number of fluorescent cells by microscopy quantification using Image-J/Fiji[2]. (A) Cells were passed through multiple subcultures without the addition of any selecting agent. Most cells continue to be fluorescent suggesting the successful continuous replication of the plasmid. A loss of fluorescent would have led to an accumulation of low/no fluorescent cells. Replication without antibiotic selection is facilitated through a toxin-antitoxin system present on the plasmid. (B) A control culture of the same cells in which multiple passing was performed as in A, but with Carbenicillin added as a selective agent (100 µg/mL) for plasmid maintenance. The similarity among the survivors shown in A and B suggests that the plasmid pMB1-A is stably maintained and expresses *gfp* in *A. baumannii*.

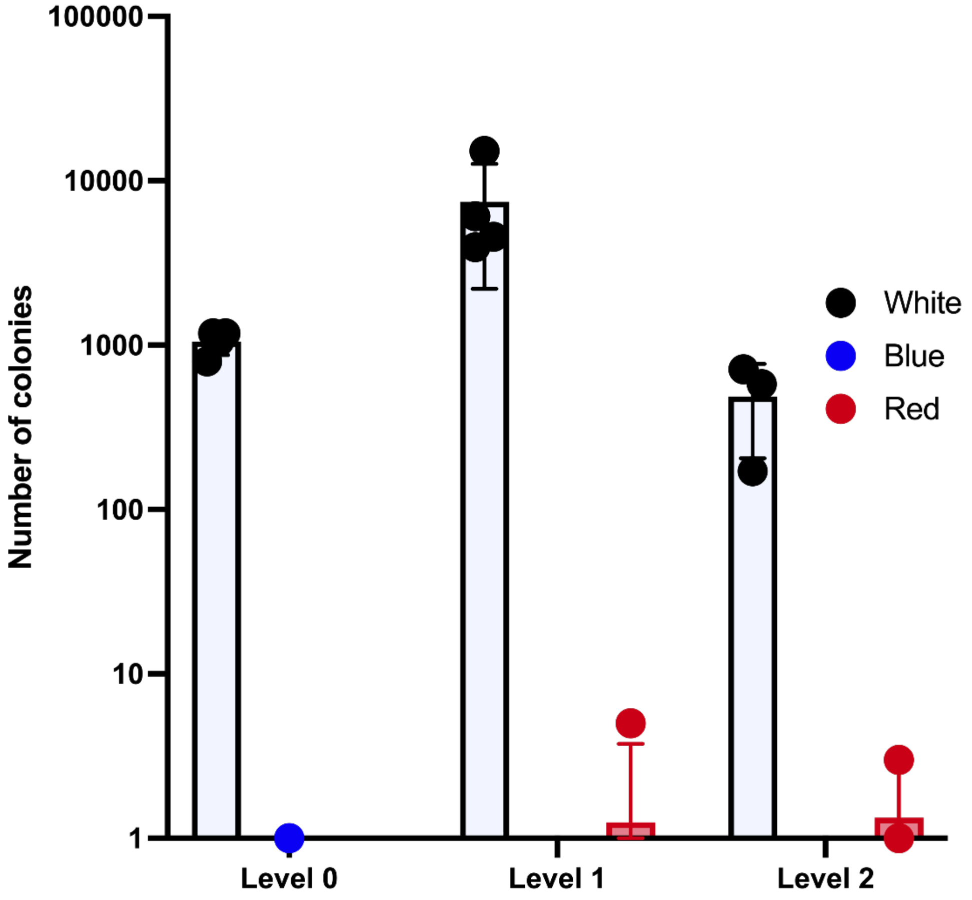


**Supplemental Figure 2**: Cloning efficiency of MoClo plasmids. The efficiency of cloning into level 0-2 was determined, and all three levels showed a remarkably high efficiency in the cloning reaction. The determination of efficiency was done by examining the color of E. coli *DH5α* that was transformed with the Golden Gate reaction mixture. The mix was deposited on LB-agar plates containing the appropriate antibiotic for each level (Chloramphenicol for level 0; Carbenicillin for level 1; kanamycin for level 2). No-insert colonies are expected to show a blue phenotype in level 0 and a red phenotype in level 1 and 2. Level 0 showed no blue phenotype colonies. Eight white colonies were sequenced, confirming the correct insertion of the desired gene. Moreover, no incorrect assembled insert was found in any of the white colonies tested.

### Supplemental Sequences

```
>pMB04(14,127bp)

ggtctcactatatctctatttaagacccactttcacatttaagttgtttttctaatccgcatatgatcaa
ttcaaggccgaataagaaggctggctctgcaccttggtgatcaaataattcgatagcttgtcgtaataat
ggcggcatactatcagtagtaggtgtttccctttcttctttagcgacttgatgctcttgatcttccaata
cgcaacctaaagtaaaatgccccacagcgctgagtgcatataatgcattctctagtgaaaaaccttgttg
gcataaaaaggctaattgattttcgagagtttcatactgtttttctgtaggccgtgtacctaaatgtact
tttgctccatcgcgatgacttagtaaagcacatctaaaacttttagcgttattacgtaaaaaatcttgcc
```

agctttccccttctaaagggcaaaagtgagtatggtgcctatctaacatctcaatggctaaggcgtcgag
caaagcccgcttatttttttacatgccaatacaatgtaggctgctctacacctagcttctgggcgagttta
cgggttgttaaaccttcgattccgacctcattaagcagctctaatgcgctgttaatcactttacttttat
ctaatctagacatcattaattcctaatttttgttgacactctatcgttgatagagttattttaccactcc
ctatcagtgatagagaaaagaattcaaaagatctgtactttaactttaagaaggagatatataaaATGGA
CTACAAAGACCATGACGGTGATTATAAAGATCATGACATCGATTACAAGGATGACGATGACAAGCTCcat
atggataagaaatactcaataggcttagctatcggcacaaatagcgtcggatgggcggtgatcactgatg
aatataaggttccgtctaaaaagttcaaggttctgggaaatacagaccgccacagtatcaaaaaaaatct
tataggggctcttttatttgacagtggagagacagcggaagcgactcgtcttaaacggacagctcgtaga
aggtatacacgtcggaagaatcgtatttgttatctacaggagattttttcaaatgagatggcgaaagtag
atgatagtttctttcatcgacttgaagagtcttttttggtggaagaagacaagaagcatgaacgtcatcc
tatttttggaaatatagtagatgaagttgcttatcatgagaaatatccaactatctatcatctgcgaaaa
aaattggtagattctactgataaagcggatttgcgcttaatctatttggccttagcgcatatgattaagt
ttcgtggtcatttttttgattgagggagatttaaatcctgataatagtgatgtggacaaactatttatcca
gttggtacaaacctacaatcaattatttgaagaaaaccctattaacgcaagtggagtagatgctaaagcg
attctttctgcacgattgagtaaatcaagacgattagaaaatctcattgctcagctccccggtgagaaga
aaaatggcttatttgggaatctcattgctttgtcattgggtttgacccctaattttaaatcaaattttga
tttggcagaagatgctaaattacagctttcaaaagatacttacgatgatgatttagataatttattggcg
caaattggagatcaatatgctgatttgtttttggcagctaagaatttatcagatgctattttactttcag
atatcctaagagtaaatactgaaataactaaggctcccctatcagcttcaatgattaaacgctacgatga
acatcatcaagacttgactcttttaaaagctttagttcgacaacaacttccagaaaagtataaagaaatc
ttttttgatcaatcaaaaaacggatatgcaggttatattgatgggggagctagccaagaagaattttata
aatttatcaaaccaatttttagaaaaaatggatggtactgaggaattattggtgaaactaaatcgtgaaga
tttgctgcgcaagcaacggacctttgacaacggctctattccccatcaaattcacttgggtgagctgcat
gctattttgagaagacaagaagacttttatccatttttaaaagacaatcgtgagaagattgaaaaaatct
tgacttttcgaattccttattatgttggtccattggcgcgtggcaatagtcgttttgcatggatgactcg
gaagtctgaagaaacaattaccccatggaattttgaagaagttgtcgataaaggtgcttcagctcaatca
tttattgaacgcatgacaaactttgataaaaatcttccaaatgaaaaagtactaccaaaacatagtttgc
tttatgagtattttacggtttataacgaattgacaaaggtcaaatatgttactgaaggaatgcgaaaacc
agcatttctttcaggtgaacagaagaaagccattgttgatttactcttcaaaacaaatcgaaaagtaacc
gttaagcaattaaaagaagattatttcaaaaaaatagaatgttttgatagtgttgaaatttcaggagttg
aagatagatttaatgcttcattaggtacctaccatgatttgctaaaaattattaaagataaagattttt
ggataatgaagaaaatgaagatatcttagaggatattgttttaacattgaccttatttgaagatagggag
atgattgaggaaagacttaaaacatatgctcacctctttgatgataaggtgatgaaacagcttaaacgtc
gccgttatactggttggggacgtttgtctcgaaaattgattaatggtattagggataagcaatctggcaa
aacaatattagatttttttgaaatcagatggttttgccaatcgcaattttatgcagctgatccatgatgat
agtttgacatttaaagaagacattcaaaaagcacaagtgtctggacaaggcgatagtttacatgaacata
ttgcaaatttagctggtagccctgctattaaaaaaggtattttacagactgtaaaagttgttgatgaatt
ggtcaaagtaatggggcggcataagccagaaaatatcgttattgaaatggcacgtgaaaatcagacaact
caaaagggccagaaaaattcgcgagagcgtatgaaacgaatcgaagaaggtatcaaagaattaggaagtc
agattcttaaagagcatcctgttgaaaatactcaattgcaaaatgaaaagctctatctctattatctcca
aaatggaagagacatgtatgtggaccaagaattagatattaatcgtttaagtgattatgatgtcgatgcc
attgttccacaaagtttccttaaagacgattcaatagacaataaggtcttaacgcgttctgataaaaatc
gtggtaaatcggataacgttccaagtgaagaagtagtcaaaaagatgaaaaactattggagacaacttct
aaacgccaagttaatcactcaacgtaagtttgataatttaacgaaagctgaacgtggaggtttgagtgaa
cttgataaagctggttttatcaaacgccaattggttgaaactcgccaaatcactaagcatgtggcacaaa
ttttggatagtcgcatgaatactaaatacgatgaaaatgataaacttattcgagaggttaaagtgattac
cttaaaatctaaattagtttctgacttccgaaaagatttccaattctataaagtacgtgagattaacaat

taccatcatgcccatgatgcgtatctaaatgccgtcgttggaactgctttgattaagaaatatccaaaac
ttgaatcggagtttgtctatggtgattataaagtttatgatgttcgtaaaatgattgctaagtctgagca
agaaataggcaaagcaaccgcaaaatatttcttttactctaatatcatgaacttcttcaaaacagaaatt
acacttgcaaatggagagattcgcaaacgccctctaatcgaaactaatggggaaactggagaaattgtct
gggataaagggcgagattttgccacagtgcgcaaagtattgtccatgccccaagtcaatattgtcaagaa
aacagaagtacagacaggcggattctccaaggagtcaattttaccaaaaagaaattcggacaagcttatt
gctcgtaaaaaagactgggatccaaaaaaatatggtggttttgatagtccaacggtagcttattcagtcc
tagtggttgctaaggtggaaaaagggaaatcgaagaagttaaaatccgttaaagagttactagggatcac
aattatggaaagaagttcctttgaaaaaaatccgattgacttttagaagctaaaggatataaggaagtt
aaaaaagacttaatcattaaactacctaaatatagtctttttgagttagaaaacggtcgtaaacggatgc
tggctagtgccggagaattacaaaaaggaaatgagctggctctgccaagcaaatatgtgaatttttata
tttagctagtcattatgaaaagttgaagggtagtccagaagataacgaacaaaaacaattgtttgtggag
cagcataagcattatttagatgagattattgagcaaatcagtgaattttctaagcgtgttattttagcag
atgccaatttagataaagttcttagtgcatataacaaacatagagacaaaccaatacgtgaacaagcaga
aaatattattcatttatttacgttgacgaatcttggagctcccgctgcttttaaatattttgatacaaca
attgatcgtaaacgatatacgtctacaaaagaagttttagatgccactcttatccatcaatccatcactg
gtctttatgaaacacgcattgatttgagtcagctaggaggtgactaactcgatcacactggctcaccttc
gggtgggcctttctgcgtttatatgtttgccctatcGATCTTTGACAGCTAGCTCAGTCCTAGGTATAAT
ACtagtcaagggcgaggaggataacaGTTTTAGAGCTAGAAATAGCAAGTTAAAATAAGGCTAGTCCGTT
ATCAACTTGAAAAAGTGGCACCGAGTCGGTGCTTTTTTTGAAGCTTtgttccggctatttgatggctagc
tcagtccttggtattatgctagcgtactttaactttaagaaggagatatacatatggtgagcaagggcga
ggaggataacatggccatcatcaaggagttcatgcgcttcaaggtgcacatggagggctccgtgaacggc
cacgagttcgagatcgagggcgagggcgagggccgcccctacgagggcacccagaccgccaagctgaagg
tgaccaagggtggcccccctgcccttcgcctgggacatcctgtcccctcagttcatgtacggctccaaggc
ctacgtgaagcaccccgccgacatccccgactacttgaagctgtccttccccgagggcttcaagtgggag
cgcgtgatgaacttcgaggacggcggcgtggtgaccgtgacccaggactcctccttgcaggacggcgagt
tcatctacaaggtgaagctgcgcggcaccaacttcccctccgacggccccgtaatgcagaagaagaccat
gggctgggaggcctcctccgagcggatgtaccccgaggacggcgccctgaagggcgagatcaagcagagg
ctgaagctgaaggacggcggccactacgacgctgaggtcaagaccacctacaaggccaagaagcccgtgc
agctgcccggcgcctacaacgtcaacatcaagttggacatcacctcccacaacgaggactacaccatcgt
ggaacagtacgaacgcgccgagggccgccactccaccggcggcatggacgagctgtacaagtaaggatcc
tcgatcacactggctcaccttcgggtgggcctttctgcgtttatatgttgaagctatttgatggctagct
cagtccttggtattatgctagcgtactttaactttaagaaggagatatacatatgcgtaaaggagaagaa
cttttcactggagttgtcccaattcttgttgaattagatggtgatgttaatgggcacaaattttctgtca
gtggagagggtgaaggtgatgcaacatacggaaaacttacccttaaatttatttgcactactggaaaact
acctgttccatggccaacacttgtcactactttcggttatggtgttcaatgctttgcgagatacccagat
catatgaaacagcatgactttttcaagagtgccatgcccgaaggttatgtacaggaaagaactatatttt
tcaaagatgacgggaactacaagacacgtgctgaagtcaagtttgaaggtgatacccttgttaatagaat
cgagttaaaaggtattgattttaaagaagatggaaacattcttggacacaaattggaatacaactataac
tcacacaatgtatacatcatggcagacaaacaaaagaatggaatcaaagttaacttcaaaattagacaca
acattgaagatggaagcgttcaactagcagaccattatcaacaaaatactccaattggcgatggccctgt
ccttttaccagacaaccattacctgtccacacaatctgccctttcgaaagatcccaacgaaaagagagat
cacatggtccttcttgagtttgtaacagctgctgggattacacatggcatggatgaactatacaaataat
cgatcacactggctcaccttcgggtgggcctttctgcgtttatatgttcttcctatcGATCTTTGACAGC
TAGCTCAGTCCTAGGTATAATACtagtaaaggagaagaacttttcacGTTTTAGAGCTAGAAATAGCAAG
TTAAAATAAGGCTAGTCCGTTATCAACTTGAAAAAGTGGCACCGAGTCGGTGCTTTTTTTGAAGCTTtgt
tTTAGgtacagagaccctgcaGTTGCAAGACAATATCGACCGcttgcagcttgataaagtgatcaaagag
ggacgagcagctacagcactgctaagggagatcggagagccactgccgccaccagtgaaagtgaaaacag

gttggtttagtgctgattattacggctttgaggacatgctcacacgttgggataatgatcaatctagcct
ttccatttatcacagagagataaaagaggagcaacaggaacagcaacgaaaacaacagcagatagaaaga
gaaactaaacaaaaacaggaagatgacaaaaagaaacggcaggactatgaaagatatttatcaaaatacg
gctatgtcaaagatgctgagtatgaccatatcagcgaccatgttgcttcagacatttccatgtttacacg
tcttcaggcgcaggcatgggcaagtaacgatatgcaagagtacataaggtgttcaaagcaaaaatacgag
cagatcagcgacagaatagcctatgagcgtgacttgaaaaaaatcgggcaattgagccgtatcctggata
aagacagacaggcacttgctgatattttaccgcagttgatgccttactatgagcagatgcaacaggctgt
taataaacggaaaattttacttgagaatgagcgacagccatcatttaaaccacgttatgaccaagaacag
cctaagccgaaaaaagacaatgacctaacttttaatgtgttacaatagcattcaaatgtaacacattgg
agtgaaaaacatgacagaagcgacttttacctttcgggtagaccatgacctaaagcaagaattttcaagc
cttgctaagactgttgatcggtcgggggcgcagctaatccgtgacttcatgcgtgactttgttaagaagc
aacaggaagccgcagactatgacaaatggttcaagcagcaggtacagatcggcttaaatgaagccaatgc
aggcaagttgattccacatgaggacgtaaaggcagagtttgccgctagacgtgctgcaacattggcgaaa
ctggcagcgaagaatgaagattgagtggactgaaacagctcgccaagatcgtagaaatatctatgattat
cttgaagaacgcaaccctatagcagctattgaaattgatgatttaattgaagaaaagacagatttacttg
ttgataatcgactgatgggcgcacaggcagacagaaagatactagggagttagtgatacatccgcatta
tgtggttgtatatgacatcactgatataatacggatactcagagtgctacacacatcgcaggagtggtca
tgacttactcatgtactttggattatttagtgttataaaatcctgatttataaatttttttgttaaaaa
agataaaagccccttgcaattgcttggggctttaccgtaatttatggggtacagatcttcgatactgaca
tatcggcaatcgaaagcattaaggtttgacgaccgctaatgatttcaccacaggggcttaatgtacctgt
cttaaattctaaggttttaactcgctttgtcaagcatagaccccaaaaatttagccaatgtctgtaactc
aatctgtccatgtgtgggtgatgaggtacagtgacgctagcacacatcggaaaaacgctattactagggg
aactgaacagagtagcggacgcaatgagtagtcatttaattggcggttatgagcgtgttcaggcggtgct
atcaatcgtaatcataacagtggcagcttgatacagtgatgtcatccctgatgcgaaagcgaccgaccga
cggtacatcgaatgggaatactttagggtgattttaagaatcgctctagggtgagtatttcccattcag
ctctgctccctccctctggtactttaatcaaaagcactactaaacatatgtttttaaataaaaaatattg
atatagagataatattagtaagaataattaaacaattgaatatagataaatcattgttaaataaagatta
attattaaaatgaatgtatacttatatataaatcaatgatttaaaatatttgataaagaaaacttttcaa
aaaaaatataattgagattgtgtcatttcggtcaattcttaatatgttccacgcaagttttagctatggt
gctaaacagaaatttgctgaaaaagaacttttcactgaactggttaaaatgtaagcagcctgagagccgc
caaaaattttaaaaacaaaccgccttaatcatcttcaaaaaatacctctaaaacctcaccatttgcgttt
taagacccatatttcatcctgcccttatgttcccatgctgatagctataaagtgtctgtaatcgcttcct
atgacgttctaggctgttgataacttttggaacaacgcaaaatgttaaaatccgatcatttttttaaccta
gttattttcgttacagggtaactttggtagtattatttcaatatttagttgttacaggataacaaaatat
atgttacagggtaattaatttagcgttacaggataactatagttacagggtaacgttaaatagttatcct
gtaatttaaaaatagttgcaggatagcataaatttttacaaataaattttataaaaagttatcctgtaac
taaaaacaatgttatcctgtaactatgatatactatatgggagttaaagacatgaaagacccgaacgatc
agaaaacacaggacatgctaaaagagccgccaaaaaccaatgccgaacgacaaaaagcatatcgtgaaaa
acgcaaaagccttgatagtcaacgtttggaggtgtttatagataagggtgtgtcggatatgcttgcggac
atggtgggcgcagcaggggagagccaaaaagccattttgaccgcattgattgagaaagagtataagcggc
tgtatgcagtgaaaaaatagcaagctaaaaaagtacattgattcaaaaagtaatgcagataaaaaagaaa
ccctcgatttatgagggtttttataactactaatgcctatgaaaataagtgcttatcattttctctaat
gtacccaataaaatcggggttttgttccacaagtttcttgatgtcattatcatgtttcttcagtaagtaa
gccccgatcaggatcttgcgttttgcatcgtctgaccttgcctgtttttgctctcgacctttcaccacag
cgtcagctagatcctttcttgccttttcctgacgttccaatttttccttttcttcatcgagcatttgacc
caattccatagctgtttttttgaccatttttaatgcctgtgtctcaacctattttccgaagtatgaacca
tcatcaatgtgaaaagtgttctacttcaaaaactcgacttacatcaagatccgagcgcagcgagcaaaat
caaaaaacaaagtcaaagccttattgctcttgcttttgcttttagcctcgcagagttcccgaagggcgca

cttacgcaaaatttttgctacgccaaattttgcaagtacggtcagggaaaccccgacaccccaaccgccc
aaaacttggggcggctgttaataaacaggtgatgaaaatggcaatttaccattgtgaaatgcagaacatt
tcgaggtcagatggtcgctcaatcgtggcatgtgcagcataccgagcaggcgaaaaattgtactgtgata
cgtacggaaaagagcaggactacaccaaaaaaacaggcattgaatacacccaaattttgccccaactgg
ggcaagtcctgacatgttagatcgtcaaaccctatggaatcgggtagagcaatccgaactaaaaaagaac
ggtgacatcaaacaggaggcaagattagcaaaggaagtagagatagcattgccgcatgaactggataaga
cacagcgtcaggcacttgttaccgagttgtgccagtccttagttaaagcctatggagtggcggtggacgt
agcgatccatgcccctcatgtgcatggggaaggagaaagaaacctcacgcccacataagacacaGGCGT
CAGGCAACTTGTTAACTGCAgtccggcaaaaaagggcaaggtgtcaccaccctgcccttttctttaaaa
ccgaaaagattacttcgcgttatgcaggcttcctcgctcactgactcgctgcgctcggtcgttcggctgc
ggcgagcggtatcagctcactcaaaggcggtaatacggttatccacagaatcaggggataacgcaggaaa
gaacatgtgagcaaaaggccagcaaaaggccaggaaccgtaaaaaggccgcgttgctggcgtttttccac
aggctccgcccccctgacgagcatcacaaaaatcgacgctcaagtcagaggtggcgaaacccgacaggac
tataaagataccaggcgtttccccctggaagctccctcgtgcgctctcctgttccgaccctgccgcttac
cggatacctgtccgcctttctcccttcgggaagcgtggcgctttctcatagctcacgctgtaggtatctc
agttcggtgtaggtcgttcgctccaagctgggctgtgtgcacgaaccccccgttcagcccgaccgctgcg
ccttatccggtaactatcgtcttgagtccaacccggtaagacacgacttatcgccactggcagcagccac
tggtaacaggattagcagagcgaggtatgtaggcggtgctacagagttcttgaagtggtggcctaactac
ggctacactagaagaacagtatttggtatctgcgctctgctgaagccagttaccttcggaaaaagagttg
gtagctcttgatccggcaaacaaaccaccgctggtagcggtggtttttttgtttgcaagcagcagattac
gcgcagaaaaaaaggatctcaagaagatcctttgatcttttctacggggtctgacgctcagtggaacgaa
aactcacgttaagggattttggtcatgagattatcaaaaaggatcttcacctagatccttttaaattaaa
aatgaagttttaaatcaatctaaagtatatatgagtaaacttggtctgacagctcgaggcttggattctc
accaataaaaaacgcccggcggcaaccgagcgttctgaacaaatccagatggagttctgaggtcattact
ggatctatcaacaggagtccaagcgagctcgatattCTCGAGAAGCTGGGGATCCGTTTGATTTTTAATG
GATAATGTGATATAATCTTTAAATACTGTAGAAAAGAGGAAGGAAATAATAAATGGCTAAAATGAGAATA
TCACCGGAATTGAAAAAACTGATCGAAAAATACCGCTGCGTAAAAGATACGGAAGGAATGTCTCCTGCTA
AGGTATATAAGCTGGTGGGAGAAAATGAAAACCTATATTTAAAAATGACGGACAGCCGGTATAAAGGGAC
CACCTATGATGTGGAACGGGAAAAGGACATGATGCTATGGCTGGAAGGAAAGCTGCCTGTTCCAAAGGTC
CTGCACTTTGAACGGCATGATGGCTGGAGCAATCTGCTCATGAGTGAGGCCGATGGCGTCCTTTGCTCGG
AAGAGTATGAAGATGAACAAAGCCCTGAAAAGATTATCGAGCTGTATGCGGAGTGCATCAGGCTCTTTCA
CTCCATCGACATATCGGATTGTCCCTATACGAATAGCTTAGACAGCCGCTTAGCCGAATTGGATTACTTA
CTGAATAACGATCTGGCCGATGTGGATTGCGAAAACTGGGAAGAAGACACTCCATTTAAAGATCCGCGCG
AGCTGTATGATTTTTTAAAGACGGAAAAGCCCGAAGAGGAACTTGTCTTTTCCCACGGCGACCTGGGAGA
CAGCAACATCTTTGTGAAAGATGGCAAAGTAAGTGGCTTTATTGATCTTGGGAGAAGCGGCAGGGCGGAC
AAGTGGTATGACATTGCCTTCTGCGTCCGGTCGATCAGGGAGGATATCGGGGAAGAACAGTATGTCGAGC
TATTTTTTGACTTACTGGGGATCAAGCCTGATTGGGAGAAAATAAAATATTATATTTTACTGGATGAATT
GTTTTAGGACGTCGCCGGCGGCATCAAATAAAACGAAAGGCTCAGTCGAAAGACTGGGCCTTTCGTTTTA
TCTGTTGTTTGTCGGTGAACGCTCTCCTGAGTAGGACAAATCCTCGAGaatatcaaattacgccccgccc
tgccactcatcgcagtactgttgtaattcattaagcattctgccgacatggaagccatcacaaacggcat
gatgaacctgaatcgccagcggcatcagcaccttgtcgccttgcgtataatatttgcccatggtgaaaac
gggggcgaagaagttgtccatattggccacgtttaaatcaaaactggtgaaactcacccagggattggct
gacacgaaaaacatattctcaataaaccctttagggaaataggccaggttttcaccgtaacacgccacat
cttgcgaatatatgtgtagaaactgccggaaatcgtcgtggtattcactccagagcgatgaaaacgtttc
agtttgctcatggaaaacggtgtaacaagggtgaacactatcccatatcaccagctcaccgtctttcatt
gccatacgaaattccggatgagcattcatcaggcgggcaagaatgtgaataaaggccggataaaacttgt
gcttatttttctttacggtctttaaaaaggccgtaatatccagctgaacggtctggttataggtacattg
agcaactgactgaaatgcctcaaaatgttctttacgatgccattgggatatatcaacggtggtatatcca

gtgatttttttctccattttagcttccttagctcctgaaaatctcgataactcaaaaaatacgcccggta
gtgatcttatttcattatggtgaaagttggaacctcttacgtgcccgatcaactcgagtgccacctgacg
tctaagaaaccattattatcatgacattaacctataaaaataggcgtatcacgaggcagaatttcagata
aaaaaaatccttagctttcgctaaggatgatttctggaattcgcggccgcttctaga

>pMB05(14,127bp)

ggtctcactatatctctatttaagacccactttcacatttaagttgtttttctaatccgcatatgatcaa
ttcaaggccgaataagaaggctggctctgcaccttggtgatcaaataattcgatagcttgtcgtaataat
ggcggcatactatcagtagtaggtgtttccctttcttctttagcgacttgatgctcttgatcttccaata
cgcaacctaaagtaaaatgccccacagcgctgagtgcatataatgcattctctagtgaaaaaccttgttg
gcataaaaaggctaattgattttcgagagtttcatactgtttttctgtaggccgtgtacctaaatgtact
tttgctccatcgcgatgacttagtaaagcacatctaaaacttttagcgttattacgtaaaaaatcttgcc
agctttccccttctaaagggcaaaagtgagtatggtgcctatctaacatctcaatggctaaggcgtcgag
caaagcccgcttattttttacatgccaatacaatgtaggctgctctacacctagcttctgggcgagttta
cgggttgttaaaccttcgattccgacctcattaagcagctctaatgcgctgttaatcactttacttttat
ctaatctagacatcattaattcctaattttttgttgacactctatcgttgatagagttattttaccactcc
ctatcagtgatagagaaaagaattcaaaagatctgtactttaactttaagaaggagatatataaaATGGA
CTACAAAGACCATGACGGTGATTATAAAGATCATGACATCGATTACAAGGATGACGATGACAAGCTCcat
atggataagaaatactcaataggcttagctatcggcacaaatagcgtcggatgggcggtgatcactgatg
aatataaggttccgtctaaaaagttcaaggttctgggaaatacagaccgccacagtatcaaaaaaaatct
tataggggctcttttatttgacagtggagagacagcggaagcgactcgtcttaaacggacagctcgtaga
aggtatacacgtcggaagaatcgtatttgttatctacaggagattttttcaaatgagatggcgaaagtag
atgatagtttctttcatcgacttgaagagtcttttttggtggaagaagacaagaagcatgaacgtcatcc
tatttttggaaatatagtagatgaagttgcttatcatgagaaatatccaactatctatcatctgcgaaaa
aaattggtagattctactgataaagcggatttgcgcttaatctatttggccttagcgcatatgattaagt
ttcgtggtcatttttttgattgagggagatttaaatcctgataatagtgatgtggacaaactatttatcca
gttggtacaaacctacaatcaattatttgaagaaaaccctattaacgcaagtggagtagatgctaaagcg
attctttctgcacgattgagtaaatcaagacgattagaaaatctcattgctcagctccccggtgagaaga
aaaatggcttatttgggaatctcattgctttgtcattgggtttgacccctaattttaaatcaaattttga
tttggcagaagatgctaaattacagctttcaaaagatacttacgatgatgatttagataatttattggcg
caaattggagatcaatatgctgatttgtttttggcagctaagaatttatcagatgctattttactttcag
atatcctaagagtaaatactgaaataactaaggctcccctatcagcttcaatgattaaacgctacgatga
acatcatcaagacttgactcttttaaaagctttagttcgacaacaacttccagaaaagtataaagaaatc
ttttttgatcaatcaaaaaacggatatgcaggttatattgatgggggagctagccaagaagaattttata
aatttatcaaaccaattttagaaaaaatggatggtactgaggaattattggtgaaactaaatcgtgaaga
tttgctgcgcaagcaacggacctttgacaacggctctattccccatcaaattcacttgggtgagctgcat
gctattttgagaagacaagaagacttttatccatttttaaaagacaatcgtgagaagattgaaaaaatct
tgacttttcgaattccttattatgttggtccattggcgcgtggcaatagtcgttttgcatggatgactcg
gaagtctgaagaaacaattaccccatggaattttgaagaagttgtcgataaaggtgcttcagctcaatca
tttattgaacgcatgacaaactttgataaaaatcttccaaatgaaaaagtactaccaaaacatagtttgc
tttatgagtattttacggtttataacgaattgacaaaggtcaaatatgttactgaaggaatgcgaaaacc
agcatttctttcaggtgaacagaagaaagccattgttgatttactcttcaaaacaaatcgaaaagtaacc
gttaagcaattaaaagaagattatttcaaaaaaatagaatgttttgatagtgttgaaatttcaggagttg
aagatagatttaatgcttcattaggtacctaccatgatttgctaaaaattattaaagataaagattttt
ggataatgaagaaaatgaagatatcttagaggatattgttttaacattgaccttatttgaagatagggag
atgattgaggaaagacttaaaacatatgctcacctctttgatgataaggtgatgaaacagcttaaacgtc
gccgttatactggttggggacgtttgtctcgaaaattgattaatggtattagggataagcaatctggcaa
aacaatattagatttttttgaaatcagatggttttgccaatcgcaattttatgcagctgatccatgatgat

agtttgacatttaaagaagacattcaaaaagcacaagtgtctggacaaggcgatagtttacatgaacata
ttgcaaatttagctggtagccctgctattaaaaaaggtattttacagactgtaaaagttgttgatgaatt
ggtcaaagtaatggggcggcataagccagaaaatatcgttattgaaatggcacgtgaaaatcagacaact
caaaagggccagaaaaattcgcgagagcgtatgaaacgaatcgaagaaggtatcaaagaattaggaagtc
agattcttaaagagcatcctgttgaaaatactcaattgcaaaatgaaaagctctatctctattatctcca
aaatggaagagacatgtatgtggaccaagaattagatattaatcgtttaagtgattatgatgtcgatgcc
attgttccacaaagtttccttaaagacgattcaatagacaataaggtcttaacgcgttctgataaaaatc
gtggtaaatcggataacgttccaagtgaagaagtagtcaaaaagatgaaaaactattggagacaacttct
aaacgccaagttaatcactcaacgtaagtttgataatttaacgaaagctgaacgtggaggtttgagtgaa
cttgataaagctggttttatcaaacgccaattggttgaaactcgccaaatcactaagcatgtggcacaaa
ttttggatagtcgcatgaatactaaatacgatgaaaatgataaacttattcgagaggttaaagtgattac
cttaaaatctaaattagtttctgacttccgaaaagatttccaattctataaagtacgtgagattaacaat
taccatcatgcccatgatgcgtatctaaatgccgtcgttggaactgctttgattaagaaatatccaaaac
ttgaatcggagtttgtctatggtgattataaagtttatgatgttcgtaaaatgattgctaagtctgagca
agaaataggcaaagcaaccgcaaaatatttcttttactctaatatcatgaacttcttcaaaacagaaatt
acacttgcaaatggagagattcgcaaacgccctctaatcgaaactaatggggaaactggagaaattgtct
gggataaagggcgagattttgccacagtgcgcaaagtattgtccatgccccaagtcaatattgtcaagaa
aacagaagtacagacaggcggattctccaaggagtcaattttaccaaaaagaaattcggacaagcttatt
gctcgtaaaaaagactgggatccaaaaaaatatggtggttttgatagtccaacggtagcttattcagtcc
tagtggttgctaaggtggaaaaagggaaatcgaagaagttaaaatccgttaaagagttactagggatcac
aattatggaaagaagttcctttgaaaaaaatccgattgactttttagaagctaaaggatataaggaagtt
aaaaaagacttaatcattaaactacctaaatatagtctttttgagttagaaaacggtcgtaaacggatgc
tggctagtgccggagaattacaaaaaggaaatgagctggctctgccaagcaaatatgtgaatttttttata
tttagctagtcattatgaaaagttgaagggtagtccagaagataacgaacaaaaacaattgtttgtggag
cagcataagcattatttagatgagattattgagcaaatcagtgaattttctaagcgtgttattttagcag
atgccaatttagataaagttcttagtgcatataacaaacatagagacaaaccaatacgtgaacaagcaga
aaatattattcatttatttacgttgacgaatcttggagctcccgctgcttttaaatattttgatacaaca
attgatcgtaaacgatatacgtctacaaaagaagttttagatgccactcttatccatcaatccatcactg
gtctttatgaaacacgcattgatttgagtcagctaggaggtgactaactcgatcacactggctcaccttc
gggtgggcctttctgcgtttatatgtttgccctatcGATCTTTGACAGCTAGCTCAGTCCTAGGTATAAT
ACtagtGTACGCGCCTGGTATGGCAGGTTTTAGAGCTAGAAATAGCAAGTTAAAATAAGGCTAGTCCGTT
ATCAACTTGAAAAAGTGGCACCGAGTCGGTGCTTTTTTTGAAGCTTtgttccggctatttgatggctagc
tcagtccttggtattatgctagcgtactttaactttaagaaggagatatacatatggtgagcaagggcga
ggaggataacatggccatcatcaaggagttcatgcgcttcaaggtgcacatggagggctccgtgaacggc
cacgagttcgagatcgagggcgagggcgagggccgcccctacgagggcacccagaccgccaagctgaagg
tgaccaagggtggccccctgcccttcgcctgggacatcctgtcccctcagttcatgtacggctccaaggc
ctacgtgaagcaccccgccgacatccccgactacttgaagctgtccttccccgagggcttcaagtgggag
cgcgtgatgaacttcgaggacggcggcgtggtgaccgtgacccaggactcctccttgcaggacggcgagt
tcatctacaaggtgaagctgcgcggcaccaacttcccctccgacggccccgtaatgcagaagaagaccat
gggctgggaggcctcctccgagcggatgtaccccgaggacggcgccctgaagggcgagatcaagcagagg
ctgaagctgaaggacggcggccactacgacgctgaggtcaagaccacctacaaggccaagaagcccgtgc
agctgcccggcgcctacaacgtcaacatcaagttggacatcacctcccacaacgaggactacaccatcgt
ggaacagtacgaacgcgccgagggccgccactccaccggcggcatggacgagctgtacaagtaaggatcc
tcgatcacactggctcaccttcgggtgggcctttctgcgtttatatgttgaagctatttgatggctagct
cagtccttggtattatgctagcgtactttaactttaagaaggagatatacatatgcgtaaaggagaagaa
cttttcactggagttgtcccaattcttgttgaattagatggtgatgttaatgggcacaaattttctgtca
gtggagagggtgaaggtgatgcaacatacggaaaacttacccttaaatttatttgcactactggaaaact
acctgttccatggccaacacttgtcactactttcggttatggtgttcaatgctttgcgagatacccagat

catatgaaacagcatgactttttcaagagtgccatgcccgaaggttatgtacaggaaagaactatatttt
tcaaagatgacgggaactacaagacacgtgctgaagtcaagtttgaaggtgatacccttgttaatagaat
cgagttaaaaggtattgattttaaagaagatggaaacattcttggacacaaattggaatacaactataac
tcacacaatgtatacatcatggcagacaaacaaaagaatggaatcaaagttaacttcaaaattagacaca
acattgaagatggaagcgttcaactagcagaccattatcaacaaaatactccaattggcgatggccctgt
ccttttaccagacaaccattacctgtccacacaatctgccctttcgaaagatcccaacgaaaagagagat
cacatggtccttcttgagtttgtaacagctgctgggattacacatggcatggatgaactatacaaataat
cgatcacactggctcaccttcgggtgggcctttctgcgtttatatgttcttcctatcGATCTTTGACAGC
TAGCTCAGTCCTAGGTATAATACtagtGTACGCGCCTGGTATGGCAGGTTTTAGAGCTAGAAATAGCAAG
TTAAAATAAGGCTAGTCCGTTATCAACTTGAAAAAGTGGCACCGAGTCGGTGCTTTTTTTGAAGCTTtgt
tTTAGgtacagagaccctgcaGTTGCAAGACAATATCGACCGcttgcagcttgataaagtgatcaaagag
ggacgagcagctacagcactgctaagggagatcggagagccactgccgccaccagtgaaagtgaaaacag
gttggtttagtgctgattattacggctttgaggacatgctcacacgttgggataatgatcaatctagcct
ttccatttatcacagagagataaaagaggagcaacaggaacagcaacgaaaacaacagcagatagaaaga
gaaactaaacaaaaacaggaagatgacaaaaagaaacggcaggactatgaaagatatttatcaaaatacg
gctatgtcaaagatgctgagtatgaccatatcagcgaccatgttgcttcagacatttccatgtttacacg
tcttcaggcgcaggcatgggcaagtaacgatatgcaagagtacataaggtgttcaaagcaaaaatacgag
cagatcagcgacagaatagcctatgagcgtgacttgaaaaaaatcgggcaattgagccgtatcctggata
aagacagacaggcacttgctgatattttaccgcagttgatgccttactatgagcagatgcaacaggctgt
taataaacggaaaattttacttgagaatgagcgacagccatcatttaaaccacgttatgaccaagaacag
cctaagccgaaaaaagacaatgacctaactttttaatgtgttacaatagcattcaaatgtaacacattgg
agtgaaaaacatgacagaagcgacttttacctttcgggtagaccatgacctaaagcaagaattttcaagc
cttgctaagactgttgatcggtcgggggcgcagctaatccgtgacttcatgcgtgactttgttaagaagc
aacaggaagccgcagactatgacaaatggttcaagcagcaggtacagatcggcttaaatgaagccaatgc
aggcaagttgattccacatgaggacgtaaaggcagagtttgccgctagacgtgctgcaacattggcgaaa
ctggcagcgaagaatgaagattgagtggactgaaacagctcgccaagatcgtagaaatatctatgattat
cttgaagaacgcaaccctatagcagctattgaaattgatgatttaattgaagaaaagacagatttacttg
ttgataatcgactgatggggcgcacaggcagacagaaagatactagggagttagtgatacatccgcatta
tgtggttgtatatgacatcactgatataatacggatactcagagtgctacacacatcgcaggagtggtca
tgacttactcatgtactttggattatttagtgttataaaatcctgatttataaattttttttgttaaaaa
agataaaagccccttgcaattgcttggggctttaccgtaatttatggggtacagatcttcgatactgaca
tatcggcaatcgaaagcattaaggtttgacgaccgctaatgatttcaccacaggggcttaatgtacctgt
cttaaattctaaggttttaactcgctttgtcaagcatagaccccaaaaatttagccaatgtctgtaactc
aatctgtccatgtgtgggtgatgaggtacagtgacgctagcacacatcggaaaaacgctattactagggg
aactgaacagagtagcggacgcaatgagtagtcatttaattggcggttatgagcgtgttcaggcggtgct
atcaatcgtaatcataacagtggcagcttgatacagtgatgtcatccctgatgcgaaagcgaccgaccga
cggtacatcgaatgggaatactttagggtgattttaagaatcgctctagggtgagtatttcccattcag
ctctgctccctccctctggtactttaatcaaaagcactactaaacatatgtttttaaataaaaaatattg
atatagagataatattagtaagaataattaaacaattgaatatagataaatcattgttaaataaagatta
attattaaaatgaatgtatacttatatataaatcaatgatttaaaatatttgataaagaaaacttttcaa
aaaaaatataattgagattgtgtcatttcggtcaattcttaatatgttccacgcaagttttagctatggt
gctaaacagaaatttgctgaaaaagaacttttcactgaactggttaaaatgtaagcagcctgagagccgc
caaaaattttaaaaacaaaccgccttaatcatcttcaaaaaatacctctaaaacctcaccatttgcgttt
taagacccatatttcatcctgcccttatgttcccatgctgatagctataaagtgtctgtaatcgcttcct
atgacgttctaggctgttgataacttttggaacaacgcaaaatgttaaaatccgatcatttttttaaccta
gttattttcgttacagggtaactttggtagtattatttcaatatttagttgttacaggataacaaaatat
atgttacagggtaattaatttagcgttacaggataactatagttacagggtaacgttaaatagttatcct
gtaatttaaaaatagttgcaggatagcataaatttttacaaataaattttataaaaagttatcctgtaac

taaaaacaatgttatcctgtaactatgatatactatatgggagttaaagacatgaaagacccgaacgatc
agaaaacacaggacatgctaaaagagccgccaaaaaccaatgccgaacgacaaaaagcatatcgtgaaaa
acgcaaaagccttgatagtcaacgtttggaggtgtttatagataagggtgtgtcggatatgcttgcggac
atggtgggcgcagcaggggagagccaaaaagccattttgaccgcattgattgagaaagagtataagcggc
tgtatgcagtgaaaaaatagcaagctaaaaaagtacattgattcaaaaagtaatgcagataaaaaagaaa
ccctcgatttatgagggtttttataactactaatgcctatgaaaataagtgcttatcatttctctaat
gtacccaataaaatcggggttttgttccacaagtttcttgatgtcattatcatgtttcttcagtaagtaa
gccccgatcaggatcttgcgttttgcatcgtctgaccttgcctgtttttgctctcgacctttcaccacag
cgtcagctagatcctttcttgccttttcctgacgttccaattttcctttcttcatcgagcatttgacc
caattccatagctgtttttttgaccattttaatgcctgtgtctcaacctattttccgaagtatgaacca
tcatcaatgtgaaaagtgttctacttcaaaaactcgacttacatcaagatccgagcgcagcgagcaaaat
caaaaaacaaagtcaaagccttattgctcttgcttttgcttttagcctcgcagagttcccgaagggcgca
cttacgcaaaattttgctacgccaaattttgcaagtacggtcagggaaaccccgacaccccaaccgccc
aaaacttggggcggctgttaataaacaggtgatgaaaatggcaatttaccattgtgaaatgcagaacatt
tcgaggtcagatggtcgctcaatcgtggcatgtgcagcataccgagcaggcgaaaaattgtactgtgata
cgtacggaaaagagcaggactacaccaaaaaaacaggcattgaatacacccaaattttgccccaactgg
ggcaagtcctgacatgttagatcgtcaaaccctatggaatcgggtagagcaatccgaactaaaaaagaac
ggtgacatcaaacaggaggcaagattagcaaaggaagtagagatagcattgccgcatgaactggataaga
cacagcgtcaggcacttgttaccgagttgtgccagtccttagttaaagcctatggagtggcggtggacgt
agcgatccatgcccctcatgtgcatggggaaggagaaagaaacctcacgcccacataagacacaGGCGT
CAGGCAACTTGTTAACTGCAgtccggcaaaaaagggcaaggtgtcaccaccctgccctttttctttaaaa
ccgaaaagattacttcgcgttatgcaggcttcctcgctcactgactcgctgcgctcggtcgttcggctgc
ggcgagcggtatcagctcactcaaaggcggtaatacggttatccacagaatcaggggataacgcaggaaa
gaacatgtgagcaaaaggccagcaaaaggccaggaaccgtaaaaaggccgcgttgctggcgtttttccac
aggctccgcccccctgacgagcatcacaaaaatcgacgctcaagtcagaggtggcgaaacccgacaggac
tataaagataccaggcgtttccccctggaagctccctcgtgcgctctcctgttccgaccctgccgcttac
cggatacctgtccgcctttctcccttcgggaagcgtggcgctttctcatagctcacgctgtaggtatctc
agttcggtgtaggtcgttcgctccaagctgggctgtgtgcacgaaccccccgttcagcccgaccgctgcg
ccttatccggtaactatcgtcttgagtccaacccggtaagacacgacttatcgccactggcagcagccac
tggtaacaggattagcagagcgaggtatgtaggcggtgctacagagttcttgaagtggtggcctaactac
ggctacactagaagaacagtatttggtatctgcgctctgctgaagccagttaccttcggaaaaagagttg
gtagctcttgatccggcaaacaaaccaccgctggtagcggtggtttttttgtttgcaagcagcagattac
gcgcagaaaaaaaggatctcaagaagatcctttgatcttttctacggggtctgacgctcagtggaacgaa
aactcacgttaagggattttggtcatgagattatcaaaaaggatcttcacctagatccttttaaattaaa
aatgaagttttaaatcaatctaaagtatatatgagtaaacttggtctgacagctcgaggcttggattctc
accaataaaaaacgcccggcggcaaccgagcgttctgaacaaatccagatggagttctgaggtcattact
ggatctatcaacaggagtccaagcgagctcgatattCTCGAGAAGCTGGGGATCCGTTTGATTTTTAATG
GATAATGTGATATAATCTTTAAATACTGTAGAAAAGAGGAAGGAAATAATAAATGGCTAAAATGAGAATA
TCACCGGAATTGAAAAAACTGATCGAAAAATACCGCTGCGTAAAAGATACGGAAGGAATGTCTCCTGCTA
AGGTATATAAGCTGGTGGGAGAAAATGAAAACCTATATTTAAAAATGACGGACAGCCGGTATAAAGGGAC
CACCTATGATGTGGAACGGGAAAAGGACATGATGCTATGGCTGGAAGGAAAGCTGCCTGTTCCAAAGGTC
CTGCACTTTGAACGGCATGATGGCTGGAGCAATCTGCTCATGAGTGAGGCCGATGGCGTCCTTTGCTCGG
AAGAGTATGAAGATGAACAAAGCCCTGAAAAGATTATCGAGCTGTATGCGGAGTGCATCAGGCTCTTTCA
CTCCATCGACATATCGGATTGTCCCTATACGAATAGCTTAGACAGCCGCTTAGCCGAATTGGATTACTTA
CTGAATAACGATCTGGCCGATGTGGATTGCGAAAACTGGGAAGAAGACACTCCATTTAAAGATCCGCGCG
AGCTGTATGATTTTTTAAAGACGGAAAAGCCCGAAGAGGAACTTGTCTTTTCCCACGGCGACCTGGGAGA
CAGCAACATCTTTGTGAAAGATGGCAAAGTAAGTGGCTTTATTGATCTTGGGAGAAGCGGCAGGGCGGAC
AAGTGGTATGACATTGCCTTCTGCGTCCGGTCGATCAGGGAGGATATCGGGGAAGAACAGTATGTCGAGC

```
TATTTTTTGACTTACTGGGGATCAAGCCTGATTGGGAGAAAATAAAATATTATATTTTACTGGATGAATT
GTTTTAGGACGTCGCCGGCGGCATCAAATAAAACGAAAGGCTCAGTCGAAAGACTGGGCCTTTCGTTTTA
TCTGTTGTTTGTCGGTGAACGCTCTCCTGAGTAGGACAAATCCTCGAGaatatcaaattacgccccgccc
tgccactcatcgcagtactgttgtaattcattaagcattctgccgacatggaagccatcacaaacggcat
gatgaacctgaatcgccagcggcatcagcaccttgtcgccttgcgtataatatttgcccatggtgaaaac
gggggcgaagaagttgtccatattggccacgtttaaatcaaaactggtgaaactcacccagggattggct
gacacgaaaaacatattctcaataaaccctttagggaaataggccaggttttcaccgtaacacgccacat
cttgcgaatatatgtgtagaaactgccggaaatcgtcgtggtattcactccagagcgatgaaaacgtttc
agtttgctcatggaaaacggtgtaacaagggtgaacactatcccatatcaccagctcaccgtctttcatt
gccatacgaaattccggatgagcattcatcaggcgggcaagaatgtgaataaaggccggataaaacttgt
gcttatttttctttacggtctttaaaaaggccgtaatatccagctgaacggtctggttataggtacattg
agcaactgactgaaatgcctcaaaatgttctttacgatgccattgggatatatcaacggtggtatatcca
gtgatttttttctccattttagcttccttagctcctgaaaatctcgataactcaaaaaatacgcccggta
gtgatcttatttcattatggtgaaagttggaacctcttacgtgcccgatcaactcgagtgccacctgacg
tctaagaaaccattattatcatgacattaacctataaaaataggcgtatcacgaggcagaatttcagata
aaaaaaatccttagctttcgctaaggatgatttctggaattcgcggccgcttctaga
```


1. Macguire, A. E. *et al.* Activation of phenotypic subpopulations in response to ciprofloxacin treatment in Acinetobacter baumannii. *Mol Microbiol* **92**, 138–152 (2014).
2. Schindelin, J. *et al.* Fiji: an open-source platform for biological-image analysis. *Nature Methods 2012 9:7* **9**, 676–682 (2012).
3. Ching, C., Gozzi, K., Heinemann, B. & Godoy, V. G. Investigating the regulation of recA in the emerging pathogen Acinetobacter baumannii. *The FASEB Journal* **31**, 591.2-591.2 (2017).
